# Comparative Studies on Architectural Stratification and Woody Species Diversity in Subtropical Evergreen Broadleaf Forests Along a Latitudinal Thermal Gradient of the Ryukyu Archipelago, Japan

**DOI:** 10.1101/007625

**Authors:** S. M. Feroz, Rempei Suwa, Koh Nakamura, Akio Hagihara, Masatsugu Yokota

## Abstract

In order to compare stand structure and woody species diversity of subtropical evergreen broadleaf forests along a latitudinal thermal gradient of the Ryukyu Archipelago, tree censuses in a 750 m^2^ plot in Okinawa Island and a 400 m^2^ plot in Ishigaki Island were performed. The number of layers increased along a latitudinal thermal gradient from four in the forest of Okinawa Island to five in the forest of Ishigaki Island. The values of Shannon’s index *H*′ and Pielou’s index *J*′ tended to increase from the top layer downward in the forest of Okinawa Island, whereas in the forest of Ishigaki Island, these values tended to increase from the bottom layer upward. High woody species diversity depended on small-sized trees in the Okinawa forest, whereas it depended on large-sized trees in the Ishigaki forest. The woody species diversity is higher in the Okinawa forest (*H*′ = 4.83 bit) than in the Ishigaki forest (*H*′ = 4.36 bit). According to successively decreasing height of layers from the top downward, the value of *H*′ increased continuously from the top layer downward in the Okinawa forest. This increasing trend was different from the Ishigaki forest, where the value of *H*′ increased up to the second layer and then decreased downward. In the Okinawa forest, the expected number of species increased continuously from the top toward the bottom layer, i.e. the bottom layer contained the highest potential number of species (65). However, in the Ishigaki forest, it increased from the top to the fourth layer and then decreased to the bottom layer, i.e. the fourth layer contained the highest potential number of species (90). The floristic composition in the Okinawa forest was different from that in the Ishigaki forest in terms of similarity index, though approximately half of the species were common between them. The highest degree of similarity in floristic composition was between the second and third layers in the Okinawa forest, whereas it was between the third and bottom layers in the Ishigaki forest. The degree of similarity in floristic composition among layers was higher in the Okinawa forest than in the Ishigaki forest. Except the top and the bottom layer respectively in the forests of Okinawa Island and Ishigaki Island, the spatial distribution of trees was random in each layer. The degree of overlapping in the spatial distribution of trees among layers in these two forests suggested that trees in the upper two layers in the Ishigaki forest can catch sufficient light, while light can not penetrate easily to the lower three layers in both of the forests. As a result, almost species in the lower layers might be shade-tolerant in both of the forests. For both of the forests, mean tree weight of each layer decreased from the top downward, whereas the corresponding tree density increased from the top downward. This trend resembled the mean weight*–*density trajectory of self-thinning plant populations.

## INTRODUCTION

The coastal areas of the Western Pacific from subarctic eastern Siberia to equatorial SE Asia have forest climates with sufficient rainfall, which develops a sequence of five forest formations: subarctic evergreen conifer forests, cool-temperate deciduous broadleaf forests, warm-temperate lucidophyll forests, subtropical forests and tropical rain forests (Kira, 1991). Within the Western Pacific sequence of thermal vegetation zones, the subtropical zone whose major part is covered by dry area, only a small part including a chain of islands from Okinawa to Taiwan and South China, is sufficiently moist to allow the development of subtropical forests. Therefore, the subtropical forests in the Ryukyu Archipelago are precious from a phytogeographical viewpoint.

A well-developed evergreen broadleaf forest exists in the northern part of Okinawa Island and in the central part of Ishigaki Island of the Ryukyu Archipelago. It is ecologically important to know how floristic composition, spatial distribution of trees, multi-layering structure and woody species diversity among layers change along a latitudinal thermal gradient of the Ryukyu Archipelago.

The diversity of tree species is fundamental to total forest biodiversity, because trees provide resources and habitats for almost all other forest species (Hall & Swaine, 1976; Huston, 1994; Whitmore, 1998; Huang et al., 2003). Measures of species diversity play a central role in ecology and conservation biology. The most commonly employed measures of species diversity are the Shannon function, species richness (number of species), and evenness (the distribution of abundance among the species, sometimes known as equitability). In addition, the spatial distribution of trees has been a major source of interest for plant ecologists because of its potential role in explaining the coexistence of tree species in species-rich forests (Bunyavejchewin et al., 2003).

The degree of canopy multi-layering and the woody species diversity increase along a latitudinal thermal gradient from higher latitudes to the tropics (Hozumi, 1975; Yamakura, 1987; Kira, 1991; Ohsawa, 1995; Kimmins, 2004). The canopy multi-layering structure, i.e. architectural stratification, is an important factor in maintaining higher woody species diversity (Roberts & Gilliam, 1995; Lindgren & Sullivan, 2001). However, there is a dearth of studies reporting the effect of the architectural stratification on floristic composition, woody species diversity and spatial distribution of trees in the subtropical evergreen broadleaf forest in Okinawa Island. It is evident that there have apparently been no studies from such a point of view in the subtropical evergreen broadleaf forest in Ishigaki Island. The purposes of this study were, therefore, to distinguish architectural stratification, to quantify woody species diversity, floristic composition and spatial distributions of trees on the basis of the architectural stratification and to compare the above parameters between the forests in the northern part (Okinawa Island) and the southern part (Ishigaki Island) of the Ryukyu Archipelago.

## MATERIALS AND METHODS

### Study sites and sampling

Two study sites were respectively selected in Okinawa Island and Ishigaki Island of the Ryukyus Archipelago. One is located in a subtropical evergreen broadleaf forest at Mt. Yonaha (26°45′N and 128°10′E), the northern part of Okinawa Island. The bedrock is composed of silicate having soil pH of 4.35 (Alhamd et al., 2004; Feroz et al., 2006). The other is located in a subtropical evergreen broadleaf forest at Mt. Omoto (24°25′03″N and 124°11′17″E), the central part of Ishigaki Island. The bedrock is composed of silicate having soil pH of 4.55.

For the former site, a sampling plot of area 750 m^2^ (25 m × 30 m) was established and divided into 120 quadrats (2.5 m × 2.5 m). The slope, altitude and direction of the plot were 24.5°, 250 m above sea level and NWW, respectively. For the latter site, a sampling plot of area 400 m^2^ (20 m × 20 m) was established and divided into 64 quadrats (2.5 m × 2.5 m). The slope and altitude and direction of the plot were 21.4°, 200 m above sea level and SW, respectively. Woody plants over 0.10 m in height for the former site and all woody plants for the latter site were numbered. They were identified to species according to the nomenclature of Walker (1976). Tree height *H* (m) and stem diameter at a height of *H*/10 *D*_0.1H_ (cm) were measured.

### Climate

The climatic data for the year of 2003∼2007 were collected from Nago Meteorological Station for the northern part of Okinawa Island and Maezato Meteorological Observatory for the central part of Ishigaki Island. Mean monthly minimum temperature and mean monthly maximum temperature are respectively 16.2 ± 0.5 (SE) °C in January and 29.1 ± 0.2 (SE) °C in July in Okinawa Island, and 18.2 ± 0.4 (SE) °C in January and 29.2 ± 0.3 (SE) °C in July in Ishigaki Island. Mean annual temperature is 22.9 ± 0.3 (SE) °C in Okinawa Island and 23.9 ± 0.3 (SE) °C in Ishigaki Island. The warmth index is 214.2 ± 0.5 (SE) °C month in Okinawa Island and 227.2 ± 1.3 (SE) °C month in Ishigaki Island, which values are within the range of 180 to 240 °C month of the subtropical region defined by Kira (1977). Mean monthly rainfall in both of the islands is over 100 mm throughout the year except for 84 ± 22 (SE) mm in February in Okinawa Island and 93 ± 19 (SE) mm in December in Ishigaki Island. Mean annual rainfall is 2050 ± 182 (SE) mm yr^-1^ in Okinawa Island and 1942 ± 159 (SE) mm yr^-1^ in Ishigaki Island. Typhoons with strong winds and rains frequently strike the islands between July and October.

### Architectural stratification

The *M–w* diagram proposed by Hozumi (1975) was used to identify the multi-layering structure of the forest stands. Tree weight *w* was assumed to be proportional to *D*_0.1H_^2^ *H* and it was arranged in descending order. Average tree weight *M_n_* from the maximum tree weight *w*_1_ to the *n*th tree weight was calculated using the form of 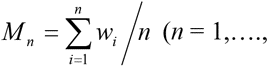 total number of trees *N*). If the *M–w* diagram is constructed by plotting the values of *M* against the corresponding values of *w* on logarithmic coordinates, then some segments on the *M–w* diagram are formed. Each segment is related to the layer with the specific characteristics of the beta–type distribution designated by Hozumi (1971, 1975). Hozumi (1975) pointed out that the segments on the *M–w* diagram can be written by either of the following equations:

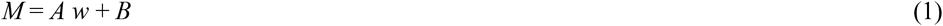

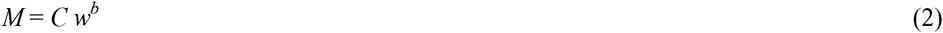

where *A*, *B*, *C* and *b* are coefficients. These functions reflect some aspect of the manners of packing trees into the three-dimensional space as realized by a forest stand.

A boundary between layers was distinguished by the relationships of the first derivative *S*_1_ and the second derivative *S*_2_ to tree weight *w* on the *M–w* diagram. The *S*_1_ and *S*_2_ are defined as follows:

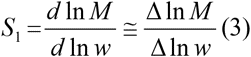

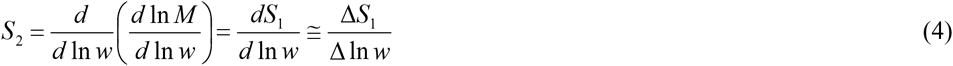

When the moving value of *S*_1_ is plotted against the corresponding value of *w*, points of inflection probably appear instead of the points of intersection between the segments on the *M–w* diagram. Then, the second derivative *S*_2_ has extrema, whose *x*-coordinates can be regarded as the points of boundary between layers.

### Species dominance

Dominance of a species was defined by importance value *IV* (%) of the species:

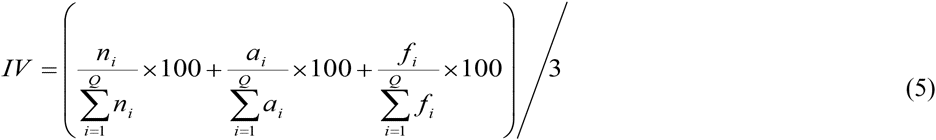

where *n_i_* is the number of individuals of the *i*th species, *a_i_* is the basal area at a height of *H*/10 of the *i*th species, *f_i_* is the number of quadrats in which the *i*th species appeared and *Q* is the total number of quadrats.

### Species-area relationship

The expected number of species *S_q_* appeared within the number of quadrats *q* selected at random from the total number of quadrats *Q* was calculated from the equation proposed by Shinozaki (1963) (cf. Hurlbert, 1971):

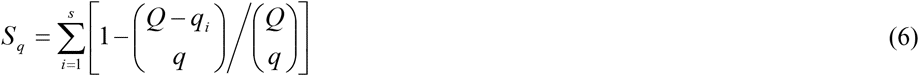

where *q_i_* is the number of quadrats in which the *i*th species occurred and *S* is the total number of species. The *S_q_*-values were obtained for *q*-values of 1, 2, 4, 8, 16, 32, 64 and 120 for the forest in Okinawa Island, and 1, 2, 4, 8, 16, 32 and 64 for the forest in Ishigaki Island.

### Floristic similarity

The similarity of floristic composition between layers was calculated using the following index *C*_Π_ (Horn, 1966; cf. Morishita, 1959; Kimoto, 1967):

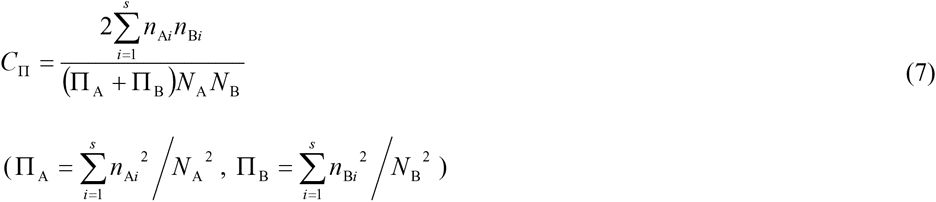

where *S* is the total number of species, *n*_A*i*_ and *n*_B*i*_ are the number of individuals of the *i*th species respectively belonging to Layer A and Layer B. The value of *C*_Π_ is 1.0 when the number of individuals belonging to a species is the same between the two layers for all species, i.e., floristic composition is completely the same between the layers, and is 0.0 when no common species is found between them.

Equation (7) was also applied for measuring the degree of similarity in floristic composition between the Okinawa and Ishigaki forests. In this case, *S* is the total number of species in the two forests, *n*_A*i*_ and *n*_B*i*_ are the number of individuals of the *i*th species respectively belonging to the Okinawa and Ishigaki forests.

### Species diversity

The following two indices of Shannon’s index (MacArthur & MacArthur, 1961) *H*′ (bit) and Pielou’s (1969) index *J*′ were used to measure woody species diversity or equitability (evenness):

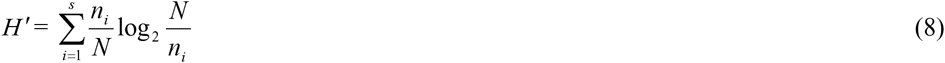

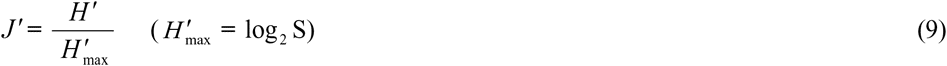

where *N* is the total number of individuals.

### Spatial distributions of trees

The unit-size 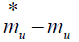 method and the *ρ*–index with successive changes of quadrat sizes (Iwao, 1972) were used to analyze the spatial distribution of trees. Mean density *m_u_* is defined as:

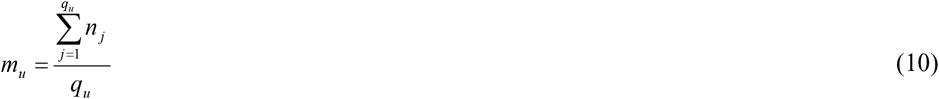

where *n*_*j*_ is the number of individuals in the *j*th quadrat and *q_u_* is the total number of quadrats when the quadrat size is *u*. The *q_u_*-values were 120, 60, 30, 15, 7, 4 and 2 respectively for the *u*-values of 1 (2.5 m × 2.5 m), 2, 4, 8, 16, 30 and 60 in the Okinawa forest, and 64, 32, 16, 8, 4 and 2 respectively for the *u*-values of 1 (2.5 m × 2.5 m), 2, 4, 8, 16 and 32 in the Ishigaki forest. On the other hand, mean crowding 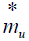 is defined by Lloyd (1967) as:

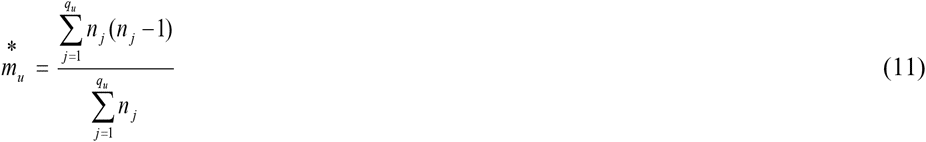

If the basic component of the spatial distribution is a single individual tree, individual trees are considered to be randomly distributed when 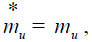 aggregately distributed when 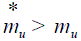 and uniformly distributed when 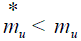 for any quadrat size. In order to provide knowledge on the distribution pattern of clumps, Iwao (1972) proposed the *ρ*–index.

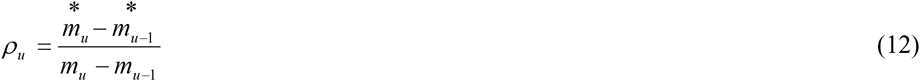

where for the smallest quadrat size (*u* = 1), 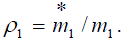 When the values of *ρ*_*u*_ are plotted against the quadrat sizes, a peak of the curve may suggest the clump area.

### Overlapping in spatial distributions of trees between layers

The *ω*-index, which is proposed by Iwao (1977) for analyzing spatial association between species, was applied to measure the degree of overlapping in spatial distributions of trees among layers with successive changes of quadrat size *u*.

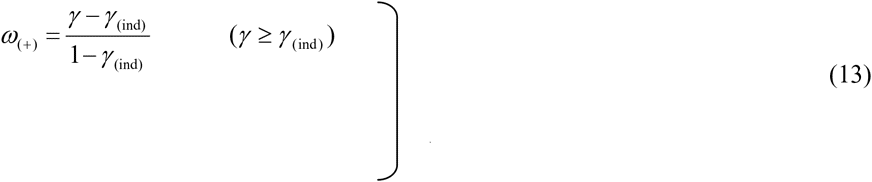

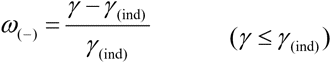

where *γ* and *γ*_(ind)_ are respectively given in the forms:

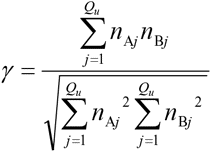

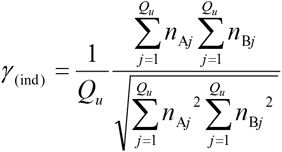

Here, *Q_u_* is the total number of quadrats taken as 120, 60, 30, 15 and 7 respectively for the *u*-values of 1 (2.5 m × 2.5 m), 2, 4, 8 and 16 in the Okinawa forest, and 64, 32, 16, 8, 4 and 2 respectively for the *u*-values of 1 (2.5 m × 2.5 m), 2, 4, 8 and 16 in the Ishigaki forest, *n*_A*j*_ and *n*_B*j*_ are the number of individuals of the *j*th quadrat respectively belonging to Layer A and Layer B. The value of *ω* changes from the maximum of + 1.0 for complete overlapping, through 0.0 for independent occurrence, to the minimum of −1.0 for complete exclusion.

### Dendrogram

The dendrograms for analyzing the degrees of floristic similarity among layers were constructed following Mountford’s (1962) method, or unweighted pair-group method using arithmetic averages (Sneath & Sokal, 1973).

### Regression analysis

The coefficients for nonlinear equations were determined with statistical analysis software (KaleidaGraph V. 4.0, Synergy Software, USA). On the other hand, the coefficients for linear and curvilinear equations were determined with the least-squares method.

## RESULTS

### Architectural stratification

The *M–w* diagram in the Okinawa forest is illustrated in Fig. 1a. As is clear from Fig. 1b, the *M–w* diagram shows four phases, the first and fourth of which phases possess a property of Eq. (1), i.e. Type I of the C*–*D curve tribe that contains eight types (Shinozaki & Kira 1961), whereas the second and third phases possess a property of Eq. (2), i.e. a power function. The *b* values in Eq. (2) were significantly different between the second and the third phases (*t* = 212, *P* ≅ 0). As a result, it was confirmed that the forest consists of four architectural layers. Figure 1c shows that the extrema, represented by arrows, emerged apparently in the *S*_2_ − ln *w* relationship. Their tree weights at boundaries between layers were estimated to be 508, 3.18 and 0.00953 cm^2^ m.

**Fig. 1.**
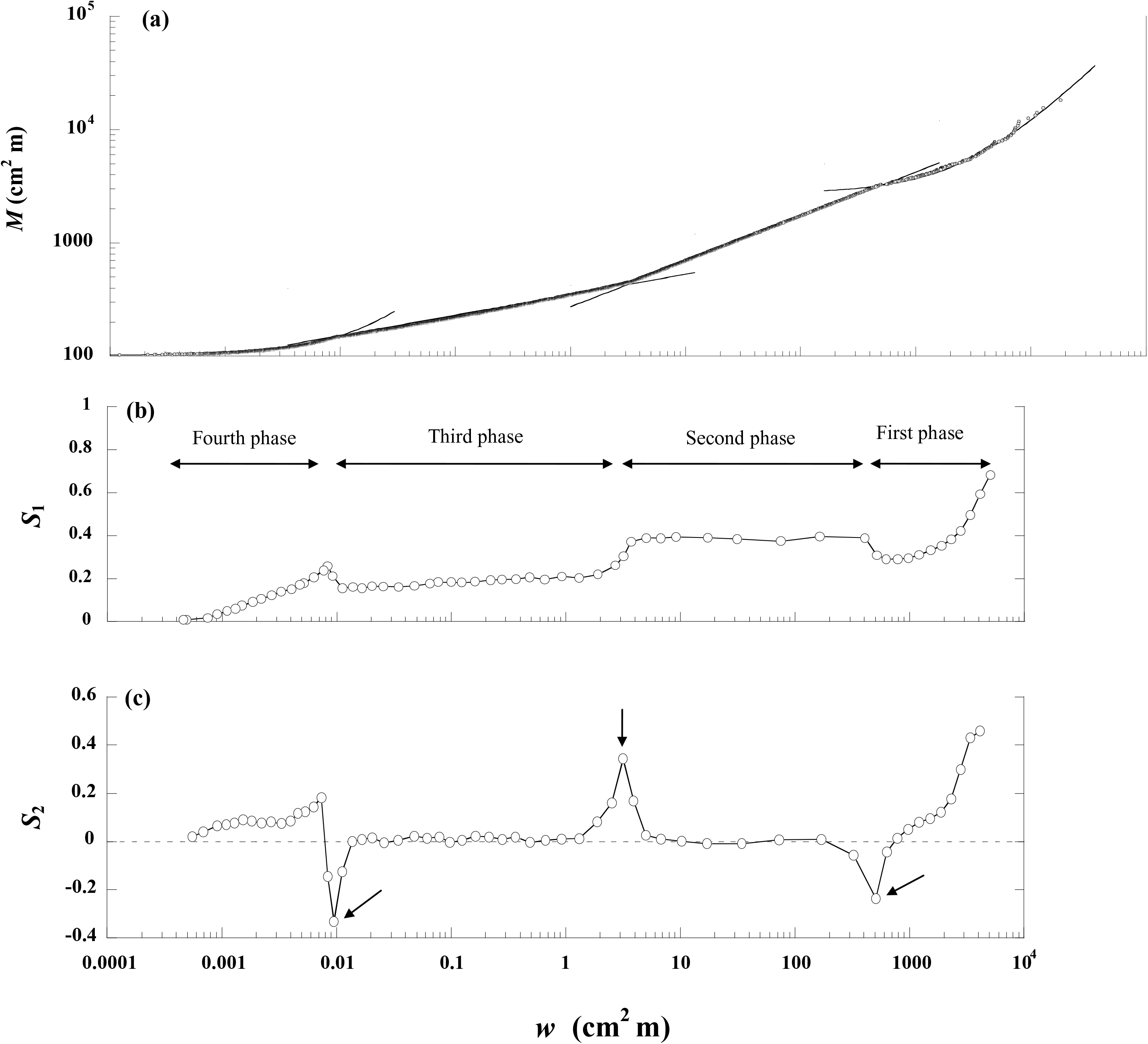
Relationships of mean tree weight *M* (**a**), the first derivative *S*_1_ (**b**) and the second derivative *S*_2_ (**c**) to tree weight *w* on logarithmic coordinates in the Okinawa forest. In the *M–w* diagram (a), the regression curves of the top and bottom layers are given by equation (1) (*R*^2^ = 0.99) for the top layer and (*R*^2^ = 0.97) for the bottom layer. The regression curves of the second and third layers are given by equation (2) (*R*^2^ = 0.91) for the second layer and (*R*^2^ = 0.94) for the third layer. The *S*_1_ and *S*_2_ are respectively defined as equations (3) and (4). The moving values of *S*_1_ and *S*_2_ are plotted against their corresponding geometric means of *w*.

On the other hand, the *M–w* in the Ishigaki forest is illustrated in Fig. 2a. As is clear from Fig. 2b, the *M–w* diagram shows five phases, all of which have the property of Eq. (1). As a result, it was confirmed that the forest consists of five architectural layers. Figure 2c shows that the extrema, represented by arrows, apparently emerged in the *S*_2_ − ln *w* relationship. Their tree weights at the boundaries between layers were estimated to be 706.3, 18.74, 0.6871 and 0.01236 cm^2^ m.

**Fig. 2.**
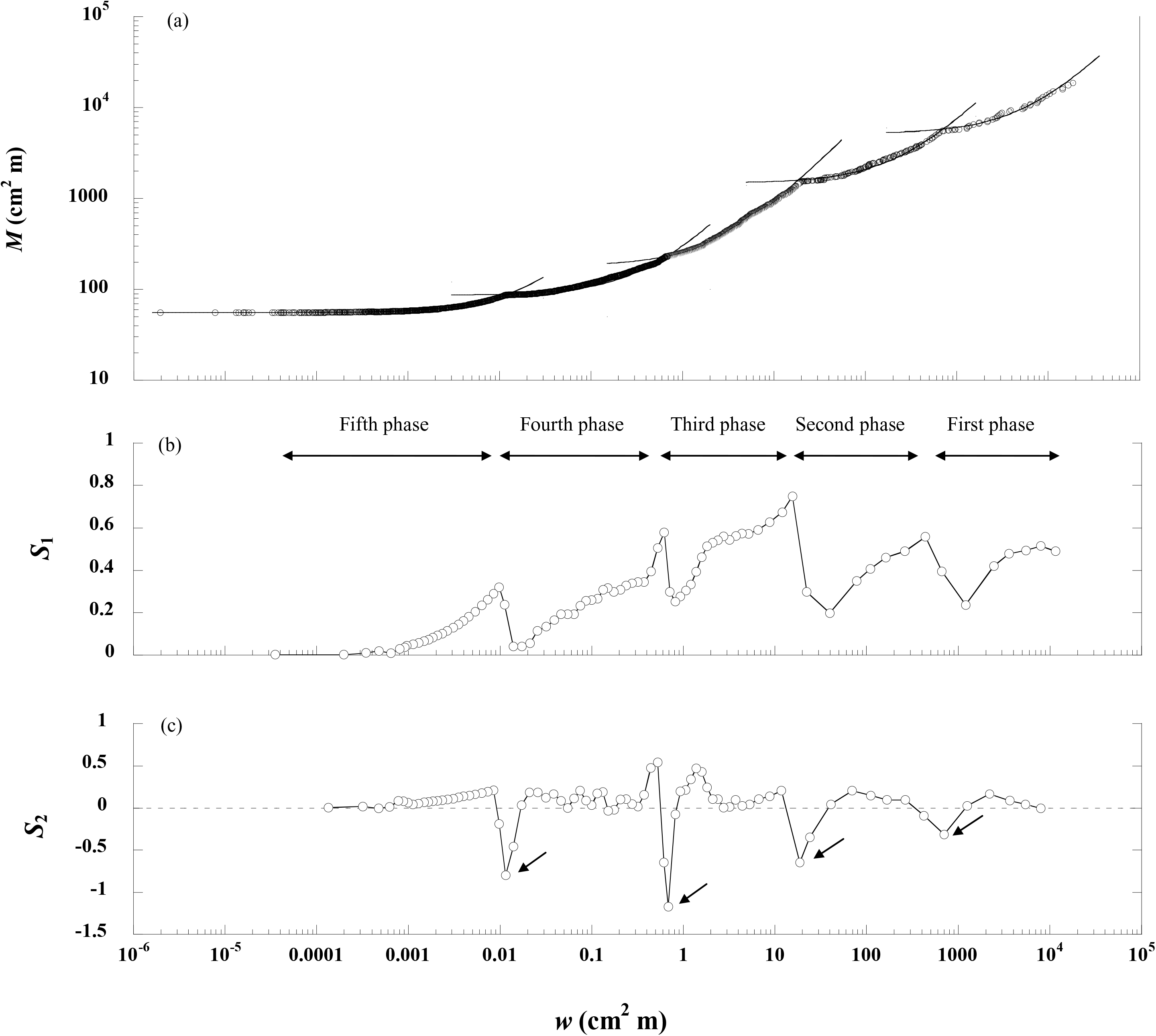
Relationships of (**a**) mean tree weight *M*, (**b**) the first derivative *S*_1_ and (**c**) the second derivative *S*_2_ to tree weight *w* on logarithmic coordinates in the Ishigaki forest. In the *M–w* diagram (a), the regression curves for all layers are given by Eq. (1) (*R*^2^ = 0.98) for the top layer, (*R*^2^ = 0.92) for the second layer, (*R*^2^ = 0.90) for the third layer, (*R*^2^ = 0.88) for the fourth layer and (*R*^2^ = 0.96) for the bottom layer. The application of *S*_1_ and *S*_2_ is the same as in Fig. 1.

Figure 3 shows the relationships between tree height *H* and weight *w* respectively for the Okinawa and Ishigaki forests, which were respectively formulated (cf. Kira & Ogawa, 1971) as follows:

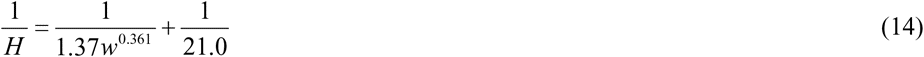

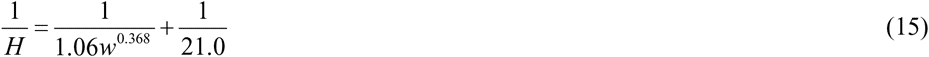

**Fig. 3.**
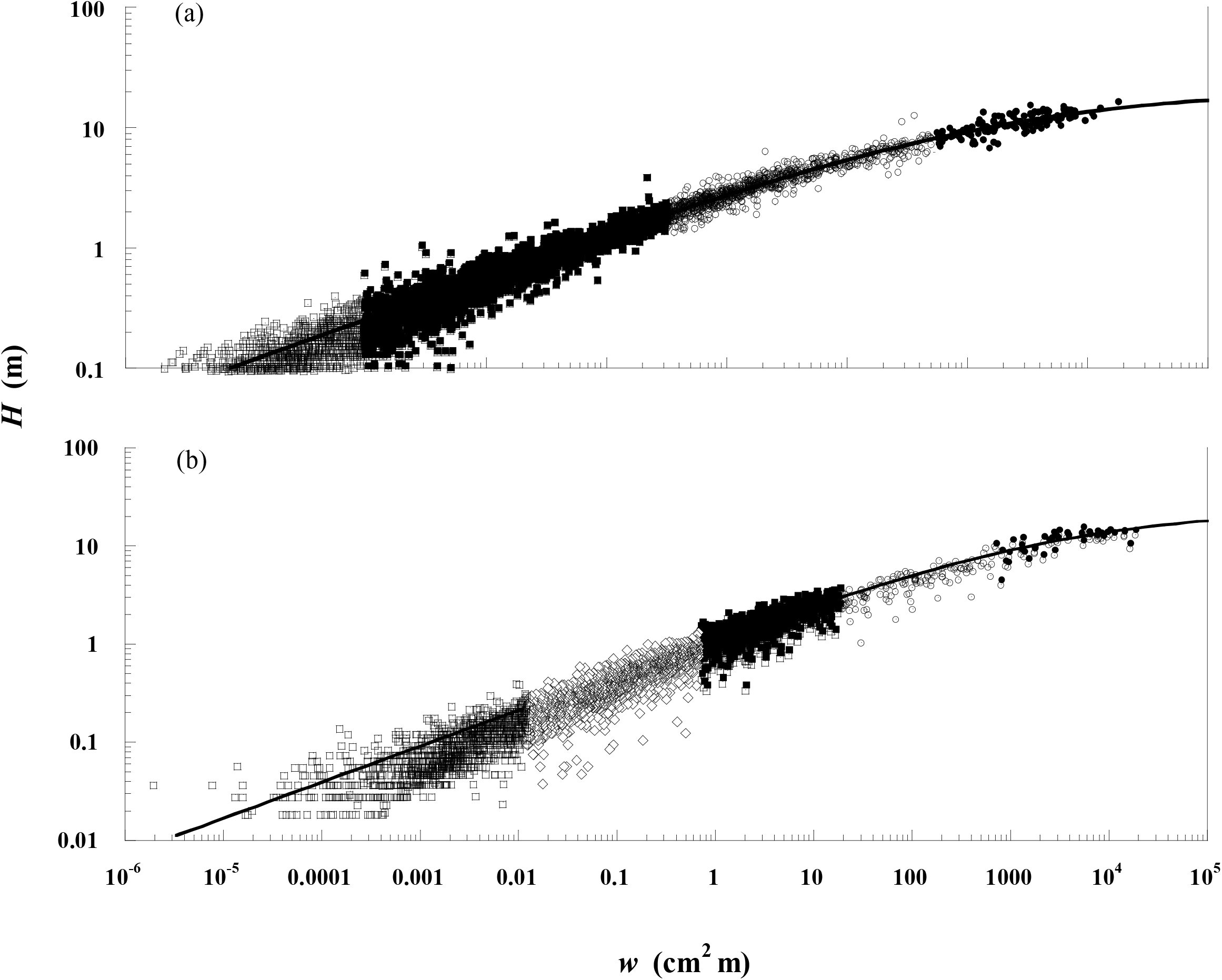
Relationships between tree height *H* and weight *w*. In the Okinawa forest **(a)**, the regression curve is given by Eq. (14) (*R*^2^ = 0.97). Filled circles, top layer (8.0 m < *H* ≤ 16.5 m); open circles, second layer (1.9 m < *H* ≤ 8.0 m); Filled squares, third layer (0.25 m < *H* ≤ 1.9 m); open squares, bottom layer (0.10 m ≤ *H* ≤ 0.25 m). In the Ishigaki forest **(b)**, the regression curve is given by Eq. (15) (*R*^2^ = 0.96). Filled circles, top layer (7.6 m < *H* ≤ 14.1 m); open circles, second layer (2.7 m < *H* ≤ 7.6 m); Filled squares, third layer (0.87 m < *H* ≤ 2.7 m); open diamond, fourth layer (0.21 m < *H* ≤ 0.87 m); open squares, bottom layer (0.0 m < *H* ≤ 0.21 m).

The estimated maximum tree height was 21.0 m, which was the same in both of the forests. The heights of the boundaries were determined as 8.0, 1.9 and 0.25 m in the Okinawa forest and 7.6, 2.7, 0.87 and 0.21 m in the Ishigaki forest by substituting the tree weights at boundaries obtained above for *w* respectively in Eqs. 14 and 15. Therefore, in the Okinawa forest, the height range was 8.0 m < *H* ≤ 16.5 m for the top layer, 1.9 m < *H* ≤ 8.0 m for the second layer, 0.25 m < *H* ≤ 1.9 m for the third layer and 0.10 m ≤ *H* ≤ 0.25 m for the bottom layer. In the Ishigaki forest, the height range was 7.6 < *H* ≤ 14.1 m for the top layer, 2.7 < *H* ≤ 7.6 m for the second layer, 0.87 < *H* ≤ 2.7 m for the third layer, 0.21 < *H* ≤ 0.87 m for the fourth layer and 0.0 < *H* ≤ 0.21 m for the bottom layer.

### Species dominance

Table 1 lists importance value *IV* of five woody species for each layer in each of the forests in order of species rank, which was determined from *IV* in the total stand. In the subtropical evergreen broadleaf forest in Okinawa Island, a total of 26 families, 43 genera, 60 species and 4684 woody individuals were recorded. The most species rich family was Rubiaceae, which contained 12 species. *Symplocos*, *Lasianthus* and *Ilex* were the species rich genera, each of which contained five species. Out of the 60 species, only three species (5%) consisted of a single individual. *Castanopsis sieboldii* (Mak.) Hatusima appeared in all layers with the highest importance value, especially with a tremendously high value of 44% in the top layer. This phenomena indicate that *C. sieboldii* is the most dominant and climax species. *Schima wallichii* (DC.) Korth. was the second dominant species in terms of *IV* of 9.30% in the total stand. A quite high *IV* of 22.8% of *S*. *wallichi* in the top layer compared with the very low *IV* (ranging from 0.62 to 1.69%) in the lower three layers indicate the heliophilic nature of the species.

**Table 1.**
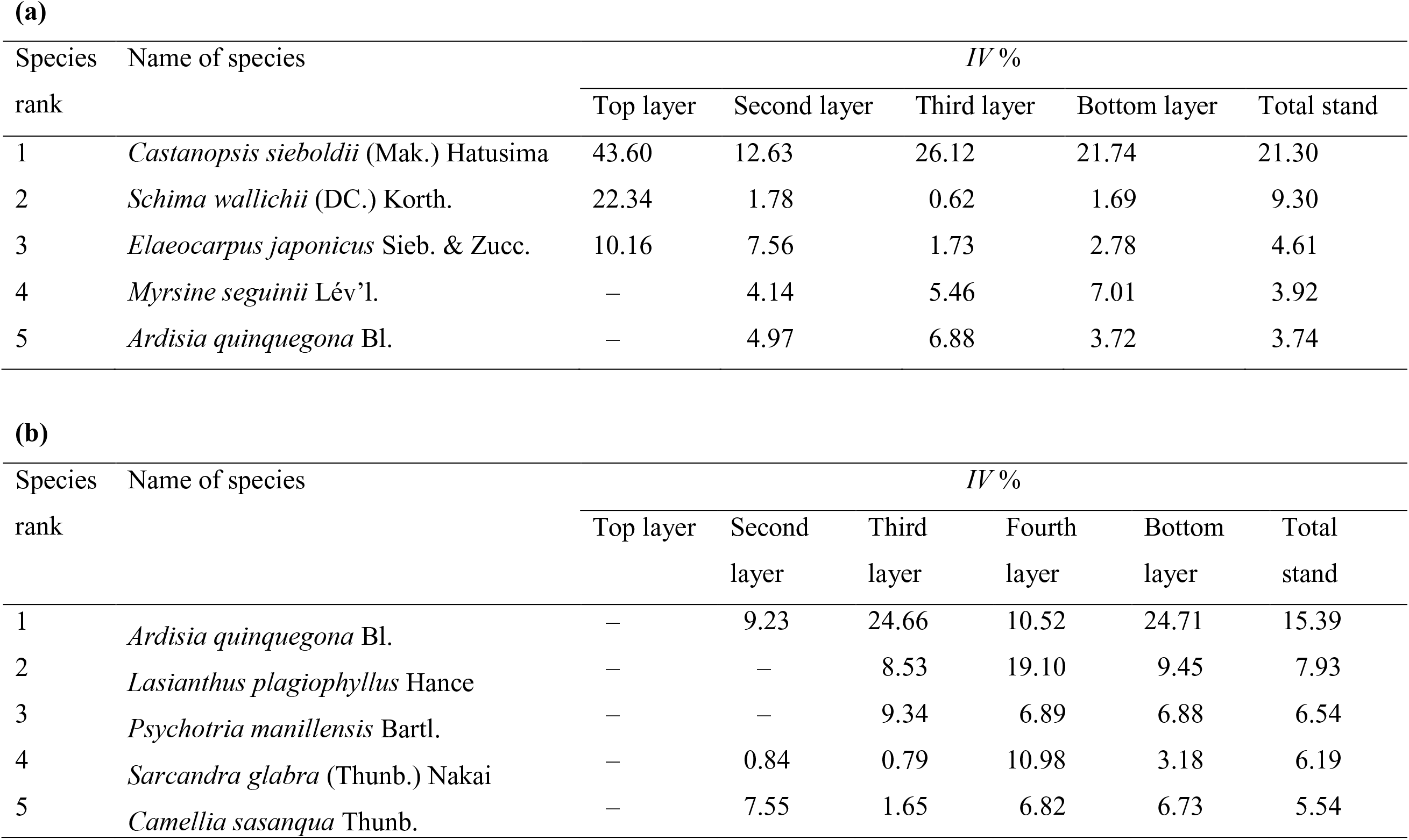
Five dominant species in the Okinawa forest **(a)** and the Ishigaki forest (**b**) in order of species rank determined by importance value *IV* in the total stand.

In the subtropical evergreen broadleaf forest in Ishigaki Island, a total of 33 families, 52 genera, 77 species and 4157 woody individuals were recorded. The most species-rich family was Aquifoliaceae, which contained six species. Aquifoliaceae consisted of *Ilex* alone, which is the most species-rich genus. Out of the 77 species, only four species (5%) consisted of a single individual. *Ardisia quinquegona* Blume was the most dominant species in terms of the highest importance value in the total stand and in all layers, except the top layer (Table 1). This species may be a subcanopy species, i.e. not a climax species, because of its disappearance in the top layer. *Castanopsis sieblodii* (Mak.) Hatusima was a low rank species in terms of the low importance value in the total stand, though it is a climax species in the Ishigaki subtropical evergreen broadleaf forest (Foster et al., 1960). However, this species appeared in all layers with a considerably higher importance value, especially the second highest importance value in the top layers (Data not shown).

### Species–area relationship

As shown in Fig. 4, the expected number of species increased, and then tended to be saturated with increasing number of quadrats for the Okinawa forest, whereas it increased with increasing number of quadrats for the Ishigaki forest. The relationships of the expected number of species *S_q_* to the number of quadrats *q* in each layer and the total stand were well approximated by the following equation (Ogawa, 1980; cf. Hagihara, 1995):

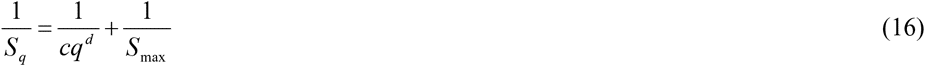

where *c* and *d* are coefficients, and *S*_max_ is the expected maximum number of species.

**Fig. 4.**
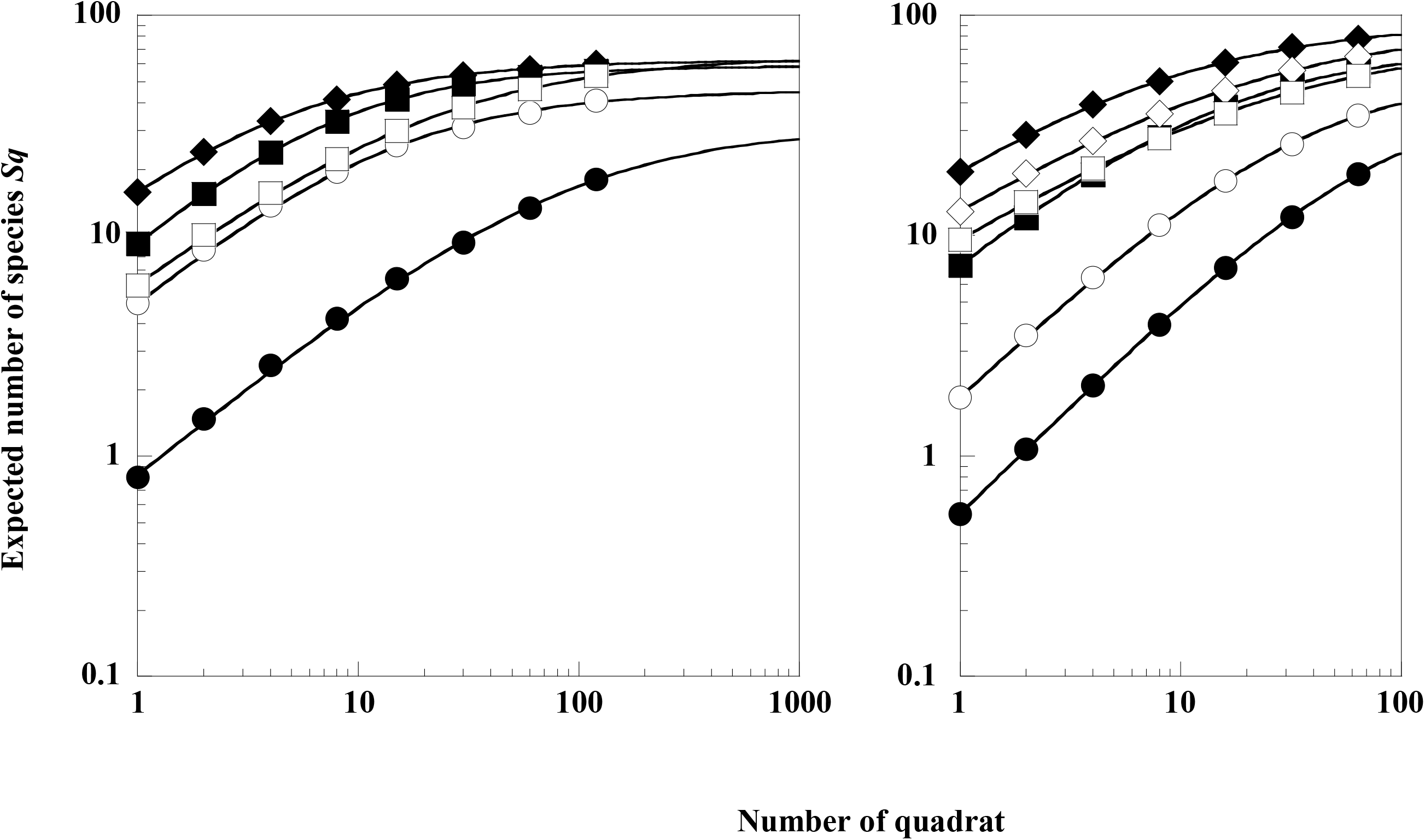
Species–area curves (area of one quadrat: 2.5 m × 2.5 m). Filled diamond, total stand and other symbols are the same as in Fig. 3. In the Okinawa forest **(a)**, the curves are given by equation (16), where *S*_max_ = 62.4 (*R*^2^ ≅ 1.0) for the total stand, *S*_max_ = 32.8 (*R*^2^ ≅ 1.0) for the top layer, *S*_max_ = 47.0 (*R*^2^ ≅ 1.0) for the second layer, *S*_max_ = 60.30 (*R*^2^ ≅ 1.0) for the third layer, *S*_max_ = 61.1 (*R*^2^ ≅ 1.0) for the bottom layer. In the Ishigaki forest **(b)**, the curves are given by the same equation, where *S*_max_ = 92.6 (*R*^2^ ≅ 1.0) for the total stand, *S*_max_ = 43.7 (*R*^2^ ≅ 1.0) for the top layer, *S*_max_ = 54.7 (*R*^2^ ≅ 1.0) for the second layer, *S*_max_ = 71.5 (*R*^2^ ≅ 1.0) for the third layer, *S*_max_ = 90.2 (*R*^2^ ≅ 1.0) for the fourth layer, *S*_max_ = 77.9 (*R*^2^ ≅ 1.0) for the bottom layer.

In the Okinawa forest, the expected maximum number of species was estimated to be 62 in the total stand. The *S*_max_ increased from 26 in the top layer, through 46 in the second layer and 61 in the third layer, to 65 in the bottom layer. Therefore, the bottom layer contained the highest potential number of species. This result was different from the Ishigaki forest, where the expected maximum number of species was estimated to be 92 in the total stand. The fourth layer contained the highest potential number of species as the *S*_max_ increased from 44 in the top layer, through 55 in the second layer and 77 in the third layer, to 90 in the fourth layer, but decreased to 88 in the bottom layer.

### Floristic similarity

The floristic similarities among layers of each forest were classified using dendrograms of similarity index *C*_Π_, as shown in Fig. 5. In the Okinawa forest, the strongest similarity in floristic composition was marked between the second and the third layers with a *C*_Π_-value of 0.80. The next was between the second-third and the bottom layers with a *C*_Π_-value of 0.64. The lowest *C*_Π_-value of 0.48 was between the top and the lower three layers. On the other hand, in the Ishigaki forest, the third and the bottom layers showed the highest similarity in floristic composition with a *C*_Π_-value of 0.94. The second and third highest similarities were respectively between the third-bottom and the fourth layers with a *C*_Π_-value of 0.69, and between the third-bottom-fourth and the second layers with a *C*_Π_-value of 0.36. The lowest *C*_Π_-value of 0.15 was between the top and the lower four layers.

**Fig. 5.**
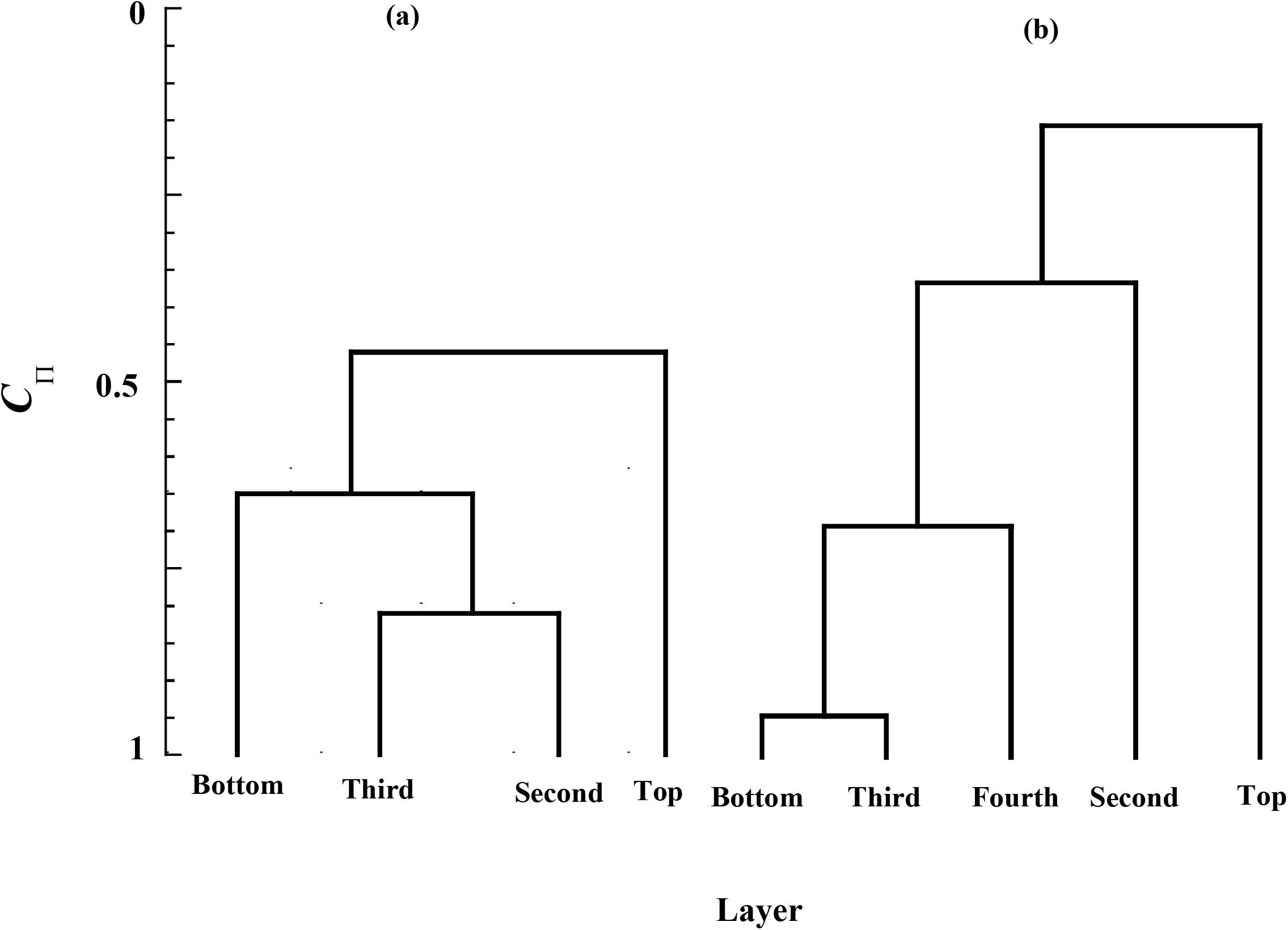
Dendrograms of the degree of similarity *C*_Π_ in floristic composition among layers in the Okinawa forest **(a)** and in the Ishigaki forest **(b).**

### Woody species diversity in the stratified forest stands

In the Okinawa forest, the values of *H*′ and *J*′ were respectively 2.75 bit and 0.66 for the top layer, 4.37 bit and 0.81 for the second layer, 4.73 bit and 0.80 for the third layer, 4.33 bit and 0.73 for the bottom layer. In the Ishigaki forest, on the other hand, the values of *H*′ and *J*′ were respectively 3.84 bit and 0.90 for the top layer, 4.49 bit and 0.87 for the second layer, 4.29 bit and 0.71 for the third layer, 4.21 bit and 0.70 for the third layer, 3.73 bit and 0.65 for the bottom layer (Table 2). These results indicate that the values of diversity indices *H*′ and *J*′ tended to increase from the top layer downward except for the bottom layer in the Okinawa forest, whereas in the Ishigaki forest they tended to increase from the bottom layer upward except for the *H*′-value of the top layer.

**Table 2.**
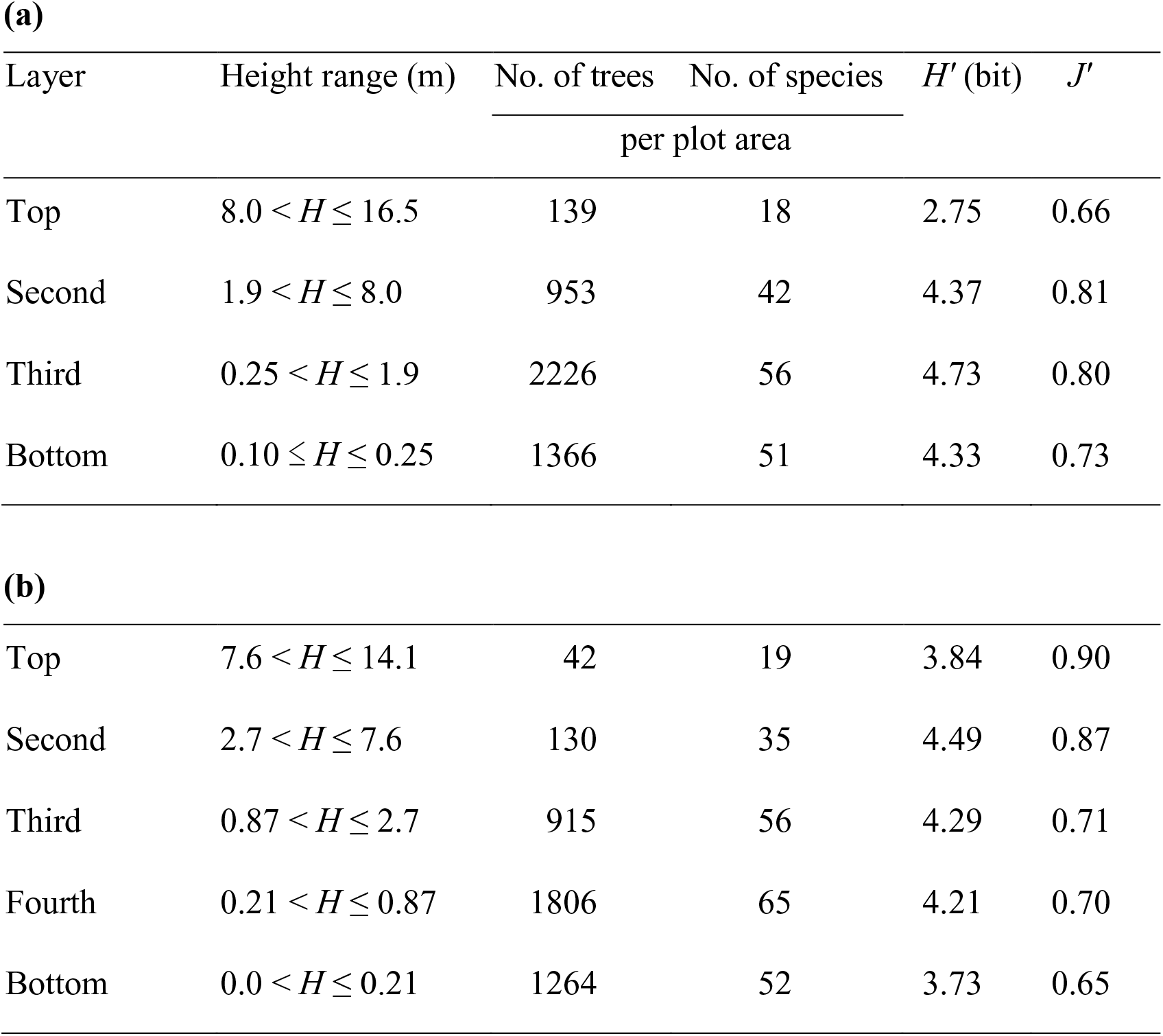
Diversity indices among layers in the Okinawa forest **(a)** and the Ishigaki forest **(b)**.

| Layer | Height range (m) | No. of trees | No. of species | $H'$ (bit) | $J'$ |
| --- | --- | --- | --- | --- | --- |
|  |  | per plot area |  |  |  |
| Top | $8.0 < H \leq 16.5$ | 139 | 18 | 2.75 | 0.66 |
| Second | $1.9 < H \leq 8.0$ | 953 | 42 | 4.37 | 0.81 |
| Third | $0.25 < H \leq 1.9$ | 2226 | 56 | 4.73 | 0.80 |
| Bottom | $0.10 \leq H \leq 0.25$ | 1366 | 51 | 4.33 | 0.73 |
| **(b)** |  |  |  |  |  |
| Top | $7.6 < H \leq 14.1$ | 42 | 19 | 3.84 | 0.90 |
| Second | $2.7 < H \leq 7.6$ | 130 | 35 | 4.49 | 0.87 |
| Third | $0.87 < H \leq 2.7$ | 915 | 56 | 4.29 | 0.71 |
| Fourth | $0.21 < H \leq 0.87$ | 1806 | 65 | 4.21 | 0.70 |
| Bottom | $0.0 < H \leq 0.21$ | 1264 | 52 | 3.73 | 0.65 |

When species diversity compared among layers, the highest value of *H*′ was in the third layer in the Okinawa forest. This is because of the second highest value of *J*′ and the highest species richness (56 species) in the third layer. However, in the Ishigaki forest, the highest value of *H*′ occurred in the second layer. This is mainly due to the second highest value of *J*′, though species richness (35 species) was the fourth highest in the second layer. On the other hand, the lowest value of *H*′ was in the top layer in the Okinawa forest. This is ascribed to the lowest *J*′-value and species richness (18 species) in the top layer. The lowest value of *H*′ was in the bottom layer. This is ascribed to the lowest value of *J*′, though species richness (52 species) was the third highest. In addition, in the Okinawa forest, the values of *H*′ in the second layer and in the bottom layer were nearly the same, while the value of *J*′ was higher in the second layer than in the bottom layer. This is because an increase of species richness from the second layer (41 species) to the bottom layer (53 species) could compensate for a decrease of evenness *J*′ from the second layer to the bottom layer. On the other hand, in the Ishigaki forest, the values of *H*′ in the top layer and in the bottom layer were nearly the same, while *J*′ was the highest in the top layer, but was the lowest in the bottom layer. This is because an increase of evenness *J*′ from the bottom layer to the top layer could compensate for a decrease of species richness from the bottom layer (52 species) to the top layer (19 species).

### Spatial distributions of trees

The spatial distribution patterns of trees for each layer are shown in Fig. 6 based on the unit-size 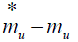 relation and the *ρ*-index against quadrat size *u*. The spatial distribution of the basic component for all layers, except the top layer in the Okinawa forest and the bottom layer in the Ishigaki forest, was a single individual and its distribution was random, because the 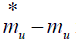 relation line mimicked the Poisson line at a significance level of 0.01. However, in the top layer of the Okinawa forest, the spatial distribution of trees showed a special trend. The *ρ*-index showed three peaks respectively at quadrat size 1, 4 and 30. The first peak at quadrat size 1 may be occasional appearance due to the underdispersion of individual trees, and the second and third peaks may perhaps be related to topography. As a result, the top layer probably consists of a double clump, the small and large clumps of which were respectively 25 m^2^ and 187.5 m^2^. In light of mean area occupied by individuals of the top layer trees in the Okinawa forest (9.1 ± 0.6 (SE) m^2^), the small clump may include three individuals and the large clump may include around eight small clumps. On the other hand, in the bottom layer of the Ishigaki forest, the spatial distribution of trees was aggregated (*P* = 1.1 × 10^−4^), though its basic component was a single individual.

**Fig. 6.**
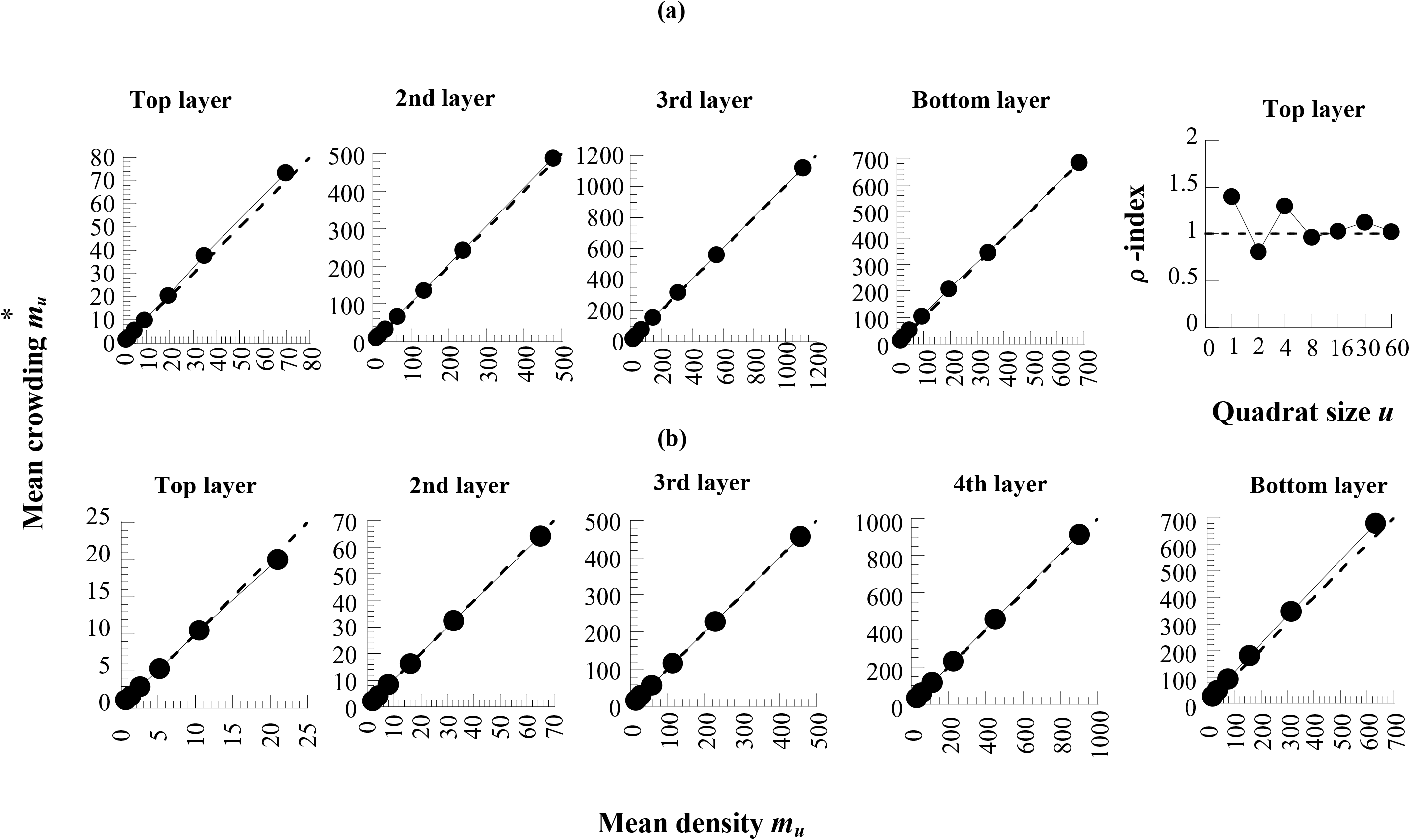
A schematic representation showing the relationships between mean crowding 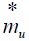 and mean density *m_u_*, and *ρ***-**index with successive changes of quadrat size *u* in the Okinawa forest **(a)** and in the Ishigaki forest **(b)**.

### Overlapping in spatial distributions of trees among layers

The degree of overlapping *ω* with successive changes of quadrat sizes in spatial distributions of trees among layers combined successively from the top layer downward is shown in Fig. 7. In the Okinawa forest, the spatial distributions of trees were overlapped between the top and the second layers, between the top-second and the third layer, and also between the top-second-third and the bottom layer. Similarly, in the Ishigaki forest, the spatial distributions of trees were nearly overlapped between the top-second and the third layers, between the top-second-third and the fourth layers, and also between the top-second-third-fourth and the bottom layers. Only a different degree of overlapping occurred between the top and the second layers, where the spatial distributions tended to be independent with each other. These results may indicate that trees in the upper two layers in the Ishigaki forest can catch sufficient light, while light can not penetrate easily to the lower three layers in both of the forests. As a result, almost species in the lower layers might be shade-tolerant in both of the forests.

**Fig. 7.**
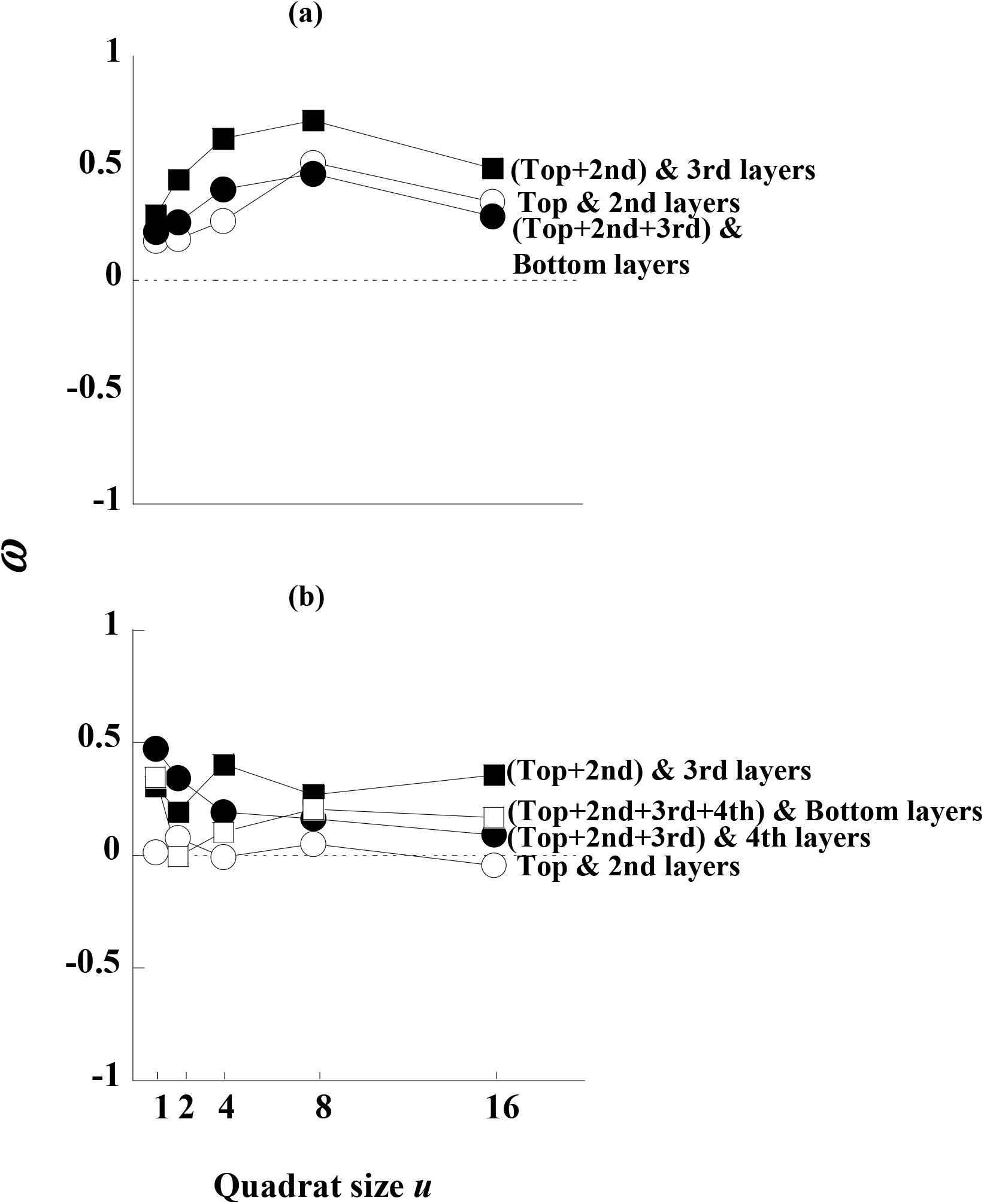
Degree of overlapping *ω* with successive changes of quadrat size *u* in spatial distributions of trees in the Okinawa forest **(a)** and in the Ishigaki forest **(b)**. The smallest quadrat size (*u* = 1) is 2.5 m × 2.5 m.

### Mean tree weight and density among layers

As shown in Fig. 8, mean tree weight 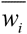 of the *i*th layer decreased from the top (*i* = 1) toward the bottom layer (*i* = 4), whereas the opposite trend was observed for the tree density *ρ_i_* of the *i*th layer. This trend was successfully expressed for the forest in Okinawa Island in the form (Feroz et al., 2006):

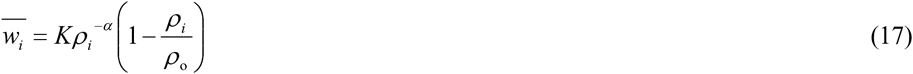

**Fig. 8.**
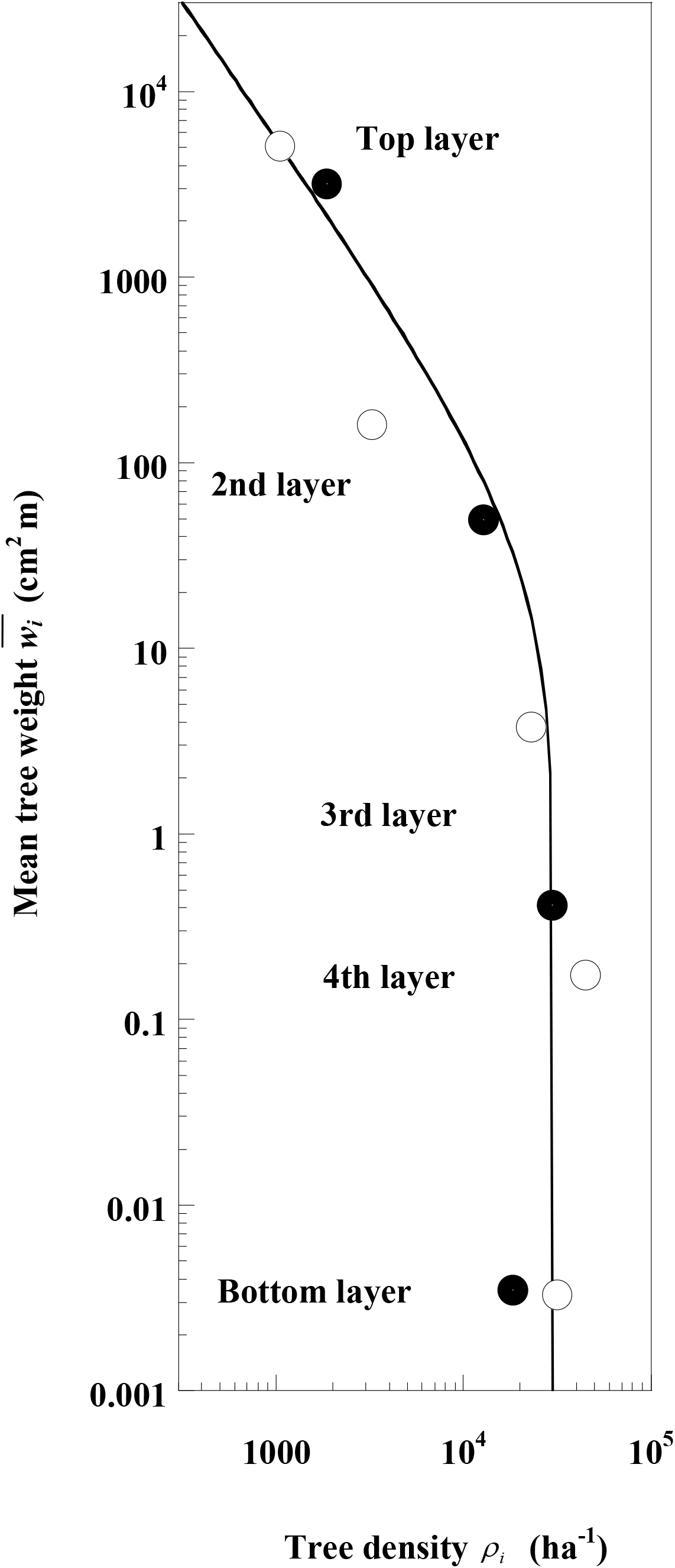
Relationship between mean tree weight 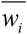 and tree density *ρ*_*i*_ among the layers. The curves are given by Eq. (17) in the Okinawa and Ishigaki forests (*R*^2^ = 0.98). Filled circle, Okinawa forest; open circle, Ishigaki forest.

In the Okinawa forest, the values of coefficients *K*, *α*, and *ρ*_o_ were estimated as 2.23 × 10^8^ cm^2^ m ha^−*α*^, 1.49 and 24575 ha^−1^. The relationship between mean tree weight 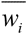 and tree density *ρ_i_* in the Ishigaki forest was well described by Eq. 15 with the same values of the coefficients obtained in the Okinawa forest (*R*^2^ = 0.98).

## DISCUSSION

All species in both of the forests in Okinawa Island and Ishigaki Island are evergreen broadleaf. The number of species in the Okinawa forest was lower than that in the Ishigaki forest, though the study area in Okinawa Island was approximately twice as large as that in Ishigaki Island. Similarly, the potential number of species in the Okinawa forest (62) was lower than that in the Ishigaki forest (92).

In both of the forests, *Castanopsis sieboldii* (Mak.) Hatusima, *Daphniphyllum teijmannii* Zoll. ex Kurz, *Neolitsea aciculata* (Bl.) Koidz. and *Dendropanax trifidus* (Thunb.) Makino were typically facultative shade species (light-tolerant under high light conditions and shade-tolerant under low light conditions) as they appeared in all layers, especially *Castanopsis sieboldii* was the most dominant species in the Okinawa forest with its highest importance value in each layer (Table 1). Although *Ardisia quinquegona* Blume was the most dominant species in the Ishigaki forest, it was not typically a facultative shade species. This is because this species did not appear in the top layer, i.e. mainly shade-tolerant, growing from the bottom layer to the subcanopy layer (or second layer). *Ardisia quinquegona* also did not appear in the top layer in the Okinawa forest, though it was one of the dominant species. *Schima wallichii* (DC.) Korth. in the Okinawa forest was typically a sun species as it appeared in the top layer with a tremendously high importance value, while it had a very low importance value in the lower three layers. In the Ishigaki forest, however, this species only appeared in the lower three layers, indicating a shade-tolerant nature which is different from the Okinawa forest.

The similarity in floristic composition between the top layer and the lower three layers in the Okinawa forest was weak, i.e. approximately one-third of the species from the lower three layers may be able to grow into the top layer. In the Ishigaki forest, however, the floristic composition between the top layer and the lower four layers was almost exclusive, i.e. approximately one-sixth of the species from the lower four layers may be able to grow into the top layer. This is because the floristic composition was more similar among layers in the Okinawa forest than in the Ishigaki forest (Fig. 5). Moreover, a low similarity in floristic composition was observed between these two forests, which is concluded from the low value of similarity index (*C*_Π_ = 0.33), though approximately half of the total number of species were common. This is mainly due to the different number of individuals belonging to the same species between these two forests. For example, the number of individuals of *Castanopsis sieboldii* (Mak.) Hatusima in the Okinawa forest (10480 ha^-1^) was not the same as in the Ishigaki forest (2600 ha^-1^).

It is known that the diversity of a community depends on two things: species richness and the evenness with which the individuals are apportioned among them (Pielou, 1975). As the lower layers contained many species relative to their smaller height ranges (Table 2), obviously these layers support high species richness of the forests. For example, 88% of the total species with 29% of the total individuals in the Okinawa forest and 67% of the total species with 30% of the total individuals in the Ishigaki forest packed within such thin bottom layers of 15 cm and 21 cm deep, respectively.

In the Okinawa forest, the value of *H*′ for small-sized trees having *H* ≥ 0.10 m was quite high as compared to that of *H*′ for large-sized trees having DBH ≥ 4.5 cm (Table 3). This is mainly caused by a large number of species for small-sized trees, though higher *J*′-value has a small influence on the high value of *H*′. As shown in Fig. 9, the trend of increasing diversity with successively decreasing height of layers from the top downward represents that high woody species diversity depended on small-sized trees. Since small-sized trees provide a natural habitat for animals living on the forest floor, conservation of small-sized trees in the lower layers is indispensable to sound maintenance of Okinawan evergreen broadleaf forests in Okinawa Island.

**Fig. 9.**
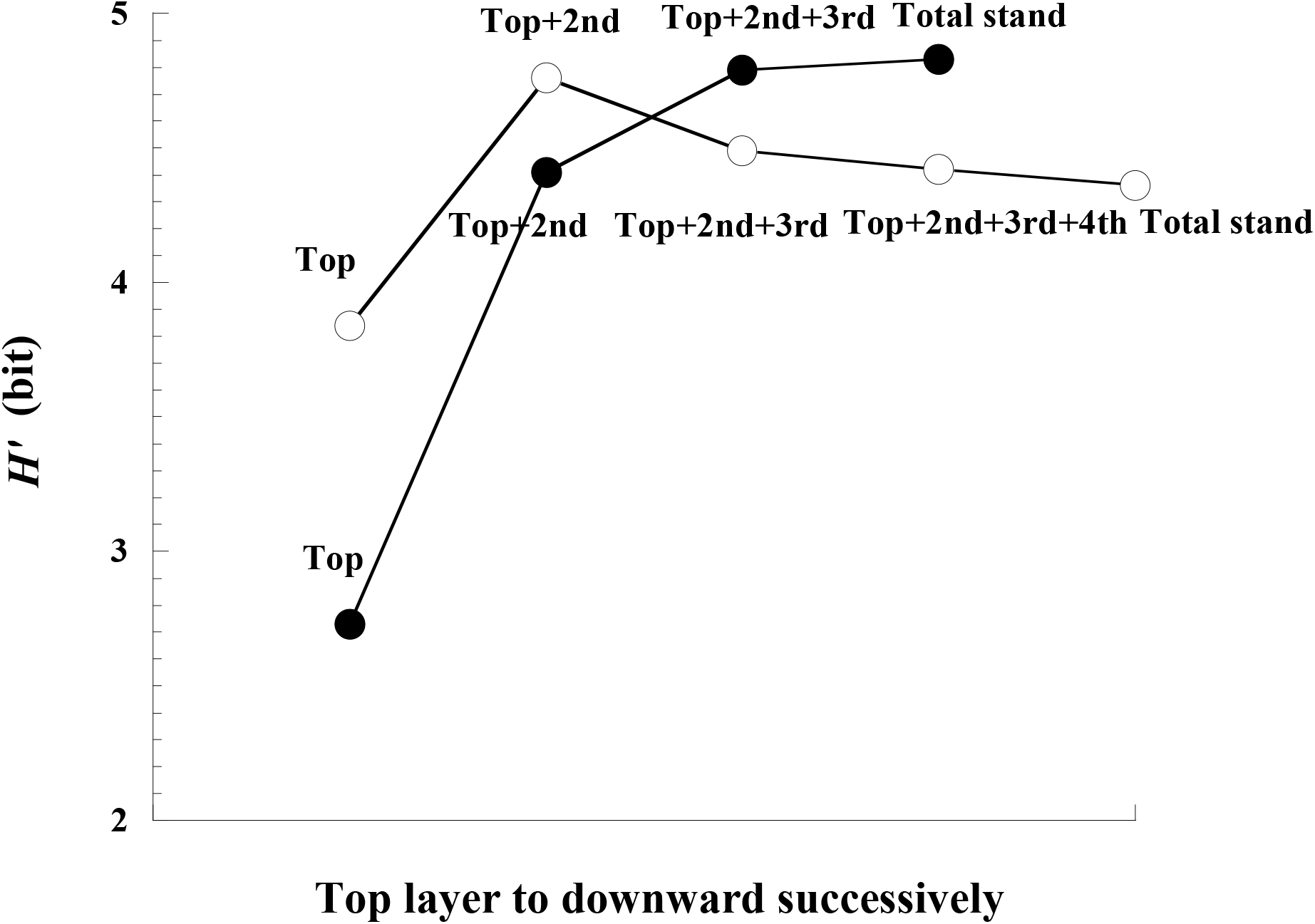
Trends of *H*′ along the layers combined from the top downward in the Okinawa forest **(a)** and in the Ishigaki forest (**b**). Symbols are the same as in Fig. 8.

**Table 3.**
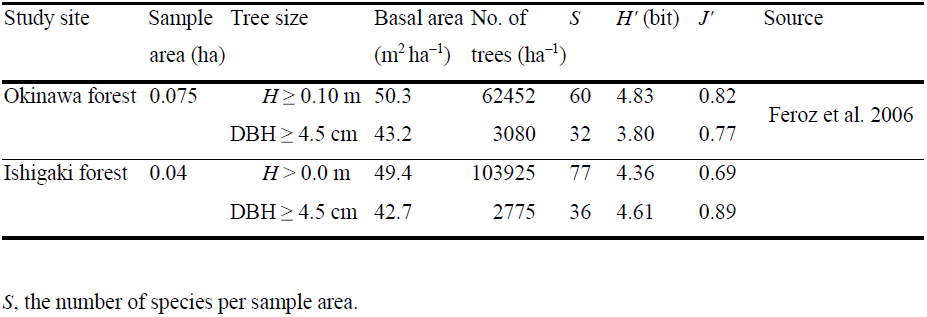
Comparison in species diversity and tree density between the Okinawa and Ishigaki forests.

On the other hand, in the Ishigaki forest, the value of *H*′ for large-sized trees having DBH ≥ 4.5 cm was quite high as compared to that of *H*′ for the total stand (Table 3). This is mainly caused by a high evenness for large-sized trees, though smaller number of species has an influence on the low value of *H*′. The diversity tends to increase up to the second layer and then decrease downward (Fig. 9), which was different from the Okinawa forest. These results may represent that the large-sized trees have an important role in maintaining high woody species diversity. Thus, it may be thought that the decreasing trend of *H*′ from the second layer downward is a characteristic of the subtropical evergreen broadleaf forest in Ishigaki Island.

The values of the diversity indices (*H*′ and *J*′) for the total stand in the Okinawa forest were higher than those in the Ishigaki forest (Table 3). This is because of a quite high evenness in the Okinawa forest compared to that in the Ishigaki forest, though species richness was lower in the Okinawa forest than in the Ishigaki forest. However, woody species diversity for large-sized trees is lower in the Okinawa forest than in the Ishigaki forest. This is caused by a higher species richness and evenness in the Okinawa forest than in the Ishigaki forest. Therefore, the statement of increasing multi-layering structure and woody species diversity along a latitudinal thermal gradient from higher latitudes to the tropics (Hozumi, 1975; Yamakura, 1987; Kira, 1991; Ohsawa, 1995; Kimmins, 2004) supported the present study for large-sized trees, but it did not support for total trees.

In Table 3, tree density of 62452 ha^−1^ for trees having *H* ≥ 0.10 m in the Okinawa forest was quite low compared to that of 103925 ha^−1^ for total trees in the Ishigaki forest, whereas basal area of 50.3 m^2^ ha^−1^ in the Okinawa forest was almost the same as that of 49.4 m^2^ ha^−1^ in the Ishigaki forest. However, tree density and basal area for trees having DBH ≥ 4.5 cm in the Okinawa forest were respectively 3080 ha^−1^and 43.2 m^2^ ha^−1^, whose values are comparable with 2775 ha^−1^and 42.7 m^2^ ha^−1^ in the Ishigaki forest. Therefore, it follows that the Ishigaki forest is densely populated with small-sized trees compared to Okinawa forest. However, the biomasses of these forest stands are possibly similar, because the basal areas are very close to each other for trees having DBH ≥ 4.5 cm.

The trend that mean tree weight decreased from the top toward the bottom layer, whereas tree density increased from the top downward (Fig. 8) was successfully formulated with Eq. 15. This trend resembles the mean plant weight*–*density trajectory of self-thinning even-aged plant populations (Hagihara, 2000), which start growing from initial plant densities lower than the initial plant density of the population obeying the −3/2 power law of self-thinning (Yoda et al., 1963). The relationship of mean tree height to tree density for the upper two layers in the Okinawa forest supported Yamakura’s quasi −1/2 power law of tree height, because the *α*-value in Eq. 15 was close to 3/2 (Feroz et al., 2006). Our formulated equation is more general than Yamakura’s quasi −1/2 power law of tree height, because the equation describes lower layers as well.

## ACKNOWLEDGEMENTS

We are grateful to our colleagues, Drs. L. Alhamd and M.N.I. Khan, Messrs. B. Tanaka, P. Wane, Sharma, K. Analuddin and Haque A.T.M. Rafiqul, and Mses. M. Wu and Y. Li, for their cooperation and active participation in the field works. Special thanks go to Prof. T. Shinzato, Subtropical Field Science Center at Yona, University of the Ryukyus, Profs. H. Ota and M. Izawa and Accos. Prof. T. Denda, Faculty of Science, University of the Ryukyus, and Prof. Z.-L. Huang, South China Institute of Botany, Chinese Academy of Science, for their kind cooperation and valuable suggestions. This study was financed in part by a Grant-in-Aid for Scientific Research (No.16201009) from the Ministry of Education, Culture, Sports, Science and Technology, and by the 21st Century COE program of the University of the Ryukyus.

